# SKiM: Accurately Classifying Metagenomic ONT Reads in Limited Memory

**DOI:** 10.1101/2025.05.13.653326

**Authors:** Trevor Schneggenburger, Jaroslaw Zola

## Abstract

**Motivation:** Oxford Nanopore Technologies’ devices, such as MinION, permit affordable, real-time DNA sequencing, and come with targeted sequencing capabilities. Such capabilities create new challenges for metagenomic classifiers that must be computationally efficient yet robust enough to handle potentially erroneous DNA reads, while ideally inspecting only a few hundred bases of a read. Currently available DNA classifiers leave room for improvement with respect to classification accuracy, memory usage, and the ability to operate in targeted sequencing scenarios.

**Results:** We present SKiM: Short K-mers in Metagenomics, a new lightweight metagenomic classifier designed for ONT reads. Compared to state-of-the-art classifiers, SKiM requires only a fraction of memory to run, and can classify DNA reads with higher accuracy after inspecting only their first few hundred bases. To achieve this, SKiM introduces new data compression techniques to maintain a reference database built from short *k*-mers, and treats classification as a statistical testing problem.

**Availability:** SKiM source code, documentation and test data are available from: https://gitlab.com/SCoRe-Group/skim.

## 1. Introduction

Oxford Nanopore Technologies (ONT) sequencing devices offer long-read DNA sequencing capabilities and can stream their sequencing data in real-time, thus allowing analysis to occur in step with sequencing. Real-time DNA analysis opens new possibilities, including mobile in-the-field deployments [8, 13, 20], as actionable information becomes available during sequencing. Such possibilities eliminate the usual delays of classic batch sequencing [26].

Real-time analysis becomes even more useful when combined with the ONT adaptive sampling feature (formerly “ReadUntil”) [11]. With adaptive sampling, a sequencer can eject a DNA molecule from a specific nanopore while it is being sequenced. After DNA ejection, the nanopore is freed up to sequence a different DNA molecule, allowing adaptive sampling to affect the content of a sequencing experiment *in silico* and improve the sequencing yield of targets of interest. So far, adaptive sampling has been used to enrich samples for low-abundance species [12], to enrich antimicrobial resistance (AMR) genes [24, 28], and to carry out metagenomic surveillance [7], to name a few applications.

However, the effectiveness of adaptive sampling depends on the analysis that decides whether or not to eject a read. In the case where a read is not of interest, it is ideal to eject it as early as possible. In this way, both the hardware and software spend minimal time processing reads that are not of interest. Therefore, this analysis must be efficient and accurate in order to handle the short starting fragments of a read.

Although the applications and benefits of real-time analysis are clear, it has some additional constraints. First, accuracy must be balanced with speed. In the worst case, the analysis stage must be able to process data at least as fast as the sequencing and basecalling, so as to not cause a bottleneck in the pipeline. Second, many of the applications of real-time analysis are done on-site. Although streaming data for off-site processing is sometimes possible, on-site applications are likely to have constrained computer hardware available, especially memory and storage. These two problems are further complicated by the rapidly increasing number of reference assemblies available [16].

Currently, real-time analysis uses one of two methods to solve the read-ejection problem. The first strategy, employed, for example, by ONT’s sequencing software (MinKNOW), is alignment of reads to a small number of reference sequences, e.g., with minimap2 [10]. While alignment is likely to produce correct matches, it is slow in practice and cannot be performed against a large number of reference sequences in real time. The second approach, provided, for example, by Readfish [19], is to depend on a traditional classifier, e.g., `Centrifuge` [6]. However, these tools are designed to work with complete reads and are computationally expensive, requiring a significant amount of RAM to run with many reference sequences. In this paper, we introduce SKiM: Short K-mers in Metagenomics. SKiM is a *k*-mer-based metagenomic classifier designed from the ground up to address the challenges of classifying ONT reads during sequencing. Compared to popular classifiers such as `Kraken2` [27] or `Centrifuge` [6], SKiM requires only a fraction of memory to run, and can classify a DNA read with high accuracy after inspecting only its first few hundred bases. Moreover, it maintains a classification throughput comparable to that of other tools. To achieve this, SKiM uses short *k*-mers (*k* = 15 or *k* = 16), data compression techniques tailored for large reference databases, and statistical correction to test significance in *k*-mers matching.

## 2. Methods

As a *k*-mer based classifier, SKiM works by decomposing both the input reads and the reference sequences into *k*-mers. What differentiates it from other *k*-mer based classifiers is how the database is stored and how *k*-mer queries are handled.

### 2.1 Database Construction

To construct a SKiM database, we begin with an input collection, *A*, of reference genome assemblies that represent classification targets. Each assembly *a*_*i*_ ∈ *A* consists of one or more DNA strings, which may correspond to chromosomes, plasmids, etc. Typically, such input data will be represented by a set of FASTA files, where each file stores a single assembly *a*_*i*_.

We convert each assembly *a*_*i*_ *∈ A* into a set, *K*_*i*_, of all *k*-mers that occur in *a*_*i*_. Importantly, each *K*_*i*_ is strictly a set, not a multiset (i.e., *K*_*i*_ stores only one copy of every found *k*-mer). We want to represent each *K*_*i*_ as a bitmap. To do so, we map each *k*- mer onto an integer in the range [0, 4^*k*^) using a bijective function *f*. While any function can be used here, in practice, *f* is such that each DNA base is assigned a 2-bit integer, and we concatenate all 2-bit integers for a *k*-mer into a 2*k*-bit integer. With this function *f*, *K*_*i*_ becomes a bitmap where *K*_*i*_[*f* (*j*)] = 1 iff the *k*-mer *j* is in the set *K*_*i*_ and 0 otherwise. Furthermore, to represent the entire collection of assemblies *A*, we combine all bitmaps into a 4^*k*^ *×* |*A*| matrix, *M*, where rows of *M* are indexed by *f* and columns are indexed by labels *K*_*i*_. *M* is the complete, uncompressed input database for SKiM. An example matrix *M* is given in Figure 1 (left). For simplicity, we use all *k*-mers in this example. In implementation, we only consider canonical *k*-mers (defined as the lexicographic minimum between a *k*-mer and its reverse complement) – a common way to account for DNA strandedness.

**Fig. 1.**
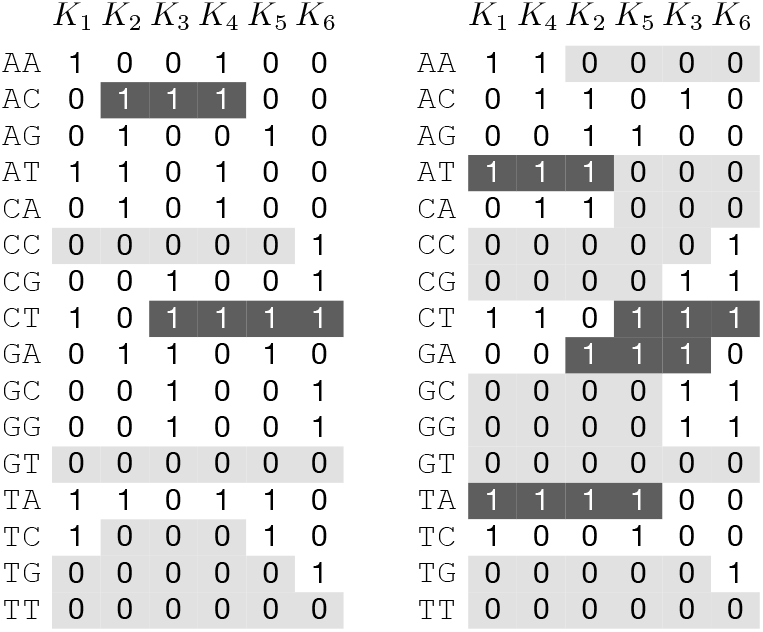
An example SKiM database with assemblies a_1_ = `AATCTAA`, a_2_ = `TAGACAT`, a_3_ = `GGACGCT`, a_4_ = `CATAACT`, a_5_ = `CTCTAGA`, a_6_ = `CCGCTGG`, and k-mers of size k = 2. Left: initial matrix M. Right: matrix M ^′^ obtained by reordering columns of M using a greedy heuristic (see Section 2.1). Runs of three or more of the same bit are highlighted.

In its basic form, matrix *M* is memory inefficient. For example, all archaeal, bacterial, and viral genomes from NCBI RefSeq, which we will refer to as ABV, currently total |*A*| ≈ 50,000 entries. Even with *k* = 15, this gives us more than 536 million distinct canonical *k*-mers that require more than 3TB of memory to store *M*.

One standard way to address this is to sub-sample the *k*-mers when forming the matrix (e.g., using minimizers [21] or winnowing [22]). In our approach, we decided to use syncmers, which have been shown to better conserve biological sequences that have errors while having a lower density than minimizers [4]. Specifically, a *k*-mer is a syncmer if its smallest *s*-mer (substring of length *s < k*) starts exactly at position *t* within the *k*-mer, where *t* is an input parameter. In our case, we chose the default values of *k* = 15, *s* = 9, and *t* = 2 (zero indexed) based on the results from [4]. This enables us to reduce the number of stored *k*-mers by approximately 85%.

While sub-sampling provides significant memory reduction, it is not sufficient on its own. For example, the sub-sampled ABV matrix would still require around 400GB to store. Hence, to further reduce the space requirements of *M*, we take advantage of two key observations. The first observation is that *M* is sparse. For instance, only about 15.5 billion bits are set in ABV, which is less than 0.5% of the total bits in the matrix. The second observation is that *M* stores the same information regardless of the ordering of the columns, provided that the columns are properly labeled by each *K*_*i*_. That is, each *K*_*i*_ can occupy an arbitrary column in *M* without affecting the way queries are processed during classification. As a consequence, we are free to decide which permutation of the columns to use. With these observations, we seek to design a compression scheme to further reduce the storage requirements of *M*.

#### Lossless Database Compression

Our goal now is to maximally compress the matrix *M* to reduce space requirements while still allowing the minimum possible theoretical runtime of queries. More specifically, the database will be queried with *k*-mers coming from DNA reads being classified (see Section 2.2). For each *k*-mer, *j*, that we query, the database should be able to retrieve all assemblies that *j* occurs in. As a lower runtime bound for such queries, consider that if *k*-mer *j* occurs in *t*_*j*_ assemblies where 0 ≤ *t*_*j*_ ≤ |*A*|, any algorithm that delivers all *t*_*j*_ assemblies must take Ω(*t*_*j*_) runtime. With this bound in mind, we consider the fact that each *k*-mer row in *M* is also a bitmap, much like each column *K*_*i*_, and has a set bit in each column (assembly) in which the *k*-mer occurs. Thus, we can compress *M* horizontally, i.e., per row (per *k*-mer), ensuring that the compression scheme allows rows to be iterable in *O*(*t*_*j*_). We note that compression in a similar context has been explored before in Taxor [25]. However, our approach differs in that we try to compress each row of *M* (that is, compression per *k*-mer), while Taxor tries to compress each column of the matrix (that is, compression per assembly). Hence, to perform the desired queries Taxor’s approach must take Ω(|*A*|) in the worst case, or on the order of the number of indexed assemblies.

We decided to use a run-length encoding (RLE) for the row-wise compression of *M*, with two main considerations in mind. First, many compression schemes, e.g., succinct data structures [14] or roaring bitmaps [3], are general purpose. This implies that auxiliary information may be stored to support lookup operations that we do not need (e.g., rank or select), which is contrary to our goal of efficiently compressing *M*. In contrast, a run-length encoding only stores the information necessary to iterate the bitmap, and it does so in the time constraints previously described. Second, there is an intuitive way to reorder columns in *M* to improve RLE compression effectiveness, which lends itself to elegant formalization. More specifically, we can try to choose an ordering of columns that creates many long runs of the same bit within the rows of *M*. Such an ordering would allow an RLE representation to encode large portions of the matrix in a small amount of space. This intuition leads to clear and well-understood criteria [5, 17] to find a column ordering of *M* such that the compression effectiveness is improved.

To explain our compression scheme, we start with the idea of a “naive” RLE (NRLE). An NRLE represents a bitmap as a list of runs, where each run corresponds to a continuous sequence of the same bit, and it encodes information about the type of bit and the number of occurrences of that bit (length of the run). For example, row CC of the left matrix in Figure 1 would consist of a run of zeros of length 5, followed by a run of ones of length 1. To achieve close to optimal compression with NRLE, we want *M* to have the consecutive ones property [1]. This property is satisfied if there exists an ordering of the columns of *M* such that all ones (that is, set bits) in each row occur consecutively. In this case, each row of *M* would require at most three runs to store, and no more than log(|*A*|) bits to encode a run. Interestingly, there exists a linear-time algorithm to test a binary matrix for the consecutive ones property using a PQ-tree data structure [1]. Unfortunately, in practice, we found that *M* is too complex to have the property even for |*A*| ≈ 10. Hence, we consider the following optimization problem instead: Let NAIVE-RUNS(*M*) be the total number of runs required to represent all rows of the matrix *M* under NRLE. We want to find a new matrix *M* ^*′*^ by permuting the columns of *M* so that naive-runs(*M* ^*′*^) is minimized.

The approach to this optimization problem has been first described in [5], and an early variant was considered in [17]. Briefly, we can imagine each column *K*_*i*_ of *M* as a point in a high-dimensional Hamming space. The Hamming distance *d*(*K*_*i*_, *K*_*j*_) between two bitmaps *K*_*i*_ and *K*_*j*_ in this space represents the number of bits that need to be substituted to edit one bitmap into the other. Imagine that we are constructing NRLEs for all rows in *M*, and we have considered bits up to column *K*_*i*_. Then, the additional number of runs needed to add a column *K*_*i*+1_ to the compression is exactly *d*(*K*_*i*_, *K*_*i*+1_), i.e., one new run for each bit that differs from *K*_*i*_ to *K*_*i*+1_. All other runs can simply be extended in length, and hence would not induce additional runs. Following this logic, naive-runs(*M*) is equivalent to the sum 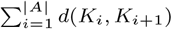, or the sum of all distances between consecutive columns in *M*. This optimization problem can then be shown to be 𝒩 *𝒫*-hard via a reduction from Hamiltonian Path [5].

Due to its computational complexity, in SKiM we solve naive-runs by treating it as an instance of the Traveling Salesperson Problem (TSP), and we apply a simple TSP greedy heuristic

(the nearest-neighbor heuristic): Starting with a random point (column), at each iteration, we add the closest point to the last added one (ignoring points that have already been added). An example result of reordering columns according to this heuristic is shown in Figure 1 (right). In this example, the reordered matrix exhibits more runs consisting of three or more bits. Moreover, we can verify that NAIVE-RUNS(*M*) = 55 while naive-runs(*M* ^*′*^) = 39.

The number of runs required to represent *M* ^*′*^ can be provably reduced further. We achieve this by adjusting our compression scheme to include uncompressed “runs” of bits when it is inefficient to represent them as runs of zeros or ones, that is, encoding their length requires more bits than they take in their uncompressed form. More specifically, let *w* be the size of a word (e.g., *w* = 16) to store a run in the RLE. A common weakness of RLEs is that very short runs from the original string, e.g., consisting of a single bit, must still be encoded using *w* bits. We bypass this limitation by allowing uncompressed “runs” within the RLE that simply store *w* bits directly as they appear in the original bitmap. We will refer to this new RLE scheme as an Adaptive RLE (ARLE). With the three types of runs (run of zeros, run of ones, uncompressed), we can use Algorithm 1 to encode an arbitrary bit string, *B*, in the fewest possible number of runs.

To compress the input bitmap *B*, the algorithm starts at the beginning of the bitmap (*i* = 0), looks for the next bit that is opposite to the one at the current position *i*. If such a bit occurs at a position *j < i* + *w*, then it is more effective to store the run *B*[*i*..*j*] uncompressed (call to the function Raw in line 9). Otherwise, the run benefits from compression as a run of zeros or ones, and it is stored in encoded form (line 12). In SKiM, we use *w* = 16 with one bit reserved to differentiate compressed and uncompressed runs, and then one more bit only for compressed runs to indicate whether it stores zeros or ones. With this, each uncompressed run represents up to *w* − 1 bits from *B*, and each compressed run encodes up to 2^*w*−2^ − 1 bits (without loss of generality, runs with more than 2^*w*−2^ − 1 bits are treated as separate runs).

Our ARLE scheme provably improves over NRLE and leads to Theorem 1 (we provide all proofs in the Supplementary Information):

**Theorem 1** *Let* ADAPTIVE-BLOCKS(*M*) *be the total number of runs required to represent any bit matrix M, obtained by running Algorithm 1 on each of the rows of M. Then* ADAPTIVE-BLOCKS(*M*) ≤ NAIVE-RUNS(*M*) *for any bit matrix M*.

We apply Theorem 1 to matrix *M* ^*′*^, i.e., we run Algorithm 1 on each row of *M* ^*′*^, to arrive at our final encoding. A summary of the complete SKiM database construction is as follows: First, each input assembly *a*_*i*_ *∈ A* is converted to a bitmap *K*_*i*_ by enumerating all sub-sampled canonical *k*-mers present in *a*_*i*_. Next, we compute the pairwise edit distances between all pairs of bitmaps (*K*_*i*_, *K*_*j*_). These distances become an input to the greedy nearest-neighbor heuristic to find an ordering of all columns to generate matrix *M* ^*′*^. With this ordering, we compute an ARLE for each row in the matrix *M* ^*′*^ according to Algorithm 1. The resulting compressed matrix is the SKiM reference database.

One final observation is that the choice of bijection *f*, which we use to order rows of *M*, has no impact on naive-runs(*M*) or ADAPTIVE-BLOCKS(*M*). However, we note that it is possible to create a partitioning, *P*, of the rows in *M* so that every set of rows *p ∈ P* can have a unique column ordering applied before ARLE. Storing the permutation of columns for each *p* would incur additional memory overhead, but by considering each partition separately, we may be able to achieve better compression. Furthermore, we could even consider dedicated variants of *f* to induce the most effective partitioning. In our initial tests, this idea introduced significant computational overhead and yielded a minimal compression improvement over the baseline.

##### Algorithm 1

Encode-as-ARLE(*B, w*)

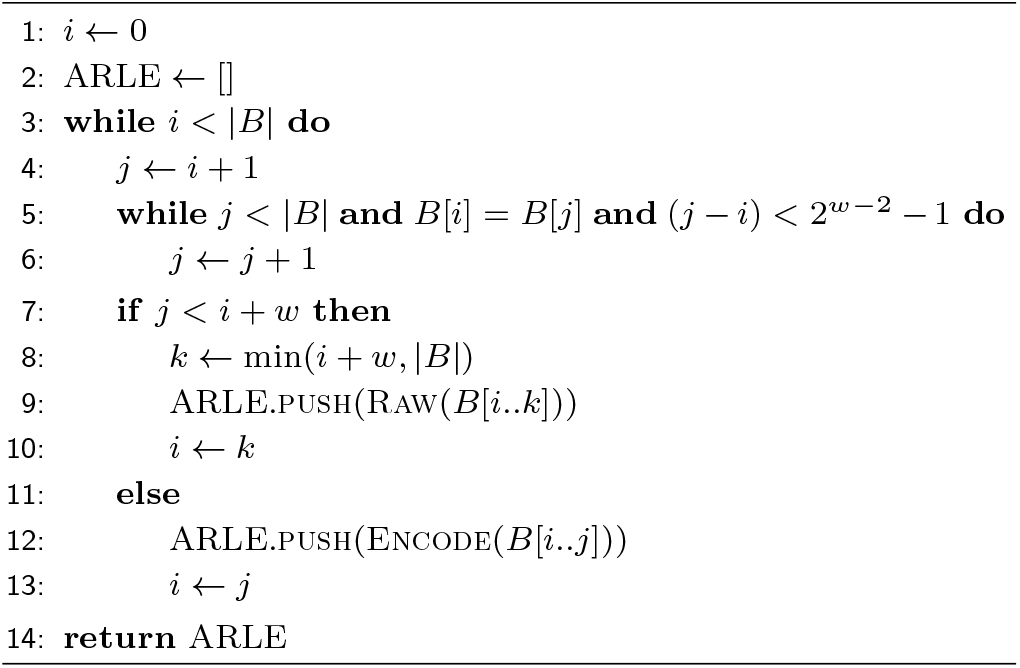

### 2.2. Classification

The use of short *k*-mers allows us to drastically reduce the size of the SKiM reference database. Moreover, in a sequence containing errors, a short *k*-mer is more likely to be error-free than a longer *k*-mer. Thus, shorter *k*-mers have an advantage during adaptive sampling, where we want to classify a sequence after inspecting only a limited number of its nucleotides. However, it also creates challenges. First, short *k*-mers are more likely to match with any reference sequence just by chance. For example, assuming our reference assembly set is *A*, if we try to classify a read that is derived from a genome not in *A*, there is a high chance that the read will contain a *k*-mer in *A* and the read will be incorrectly classified. Second, false positive *k*-mer matching may introduce a significant bias toward larger reference sequences. For example, a bacterial sequence may contain 100*×* as many *k*-mers as a viral sequence. In such a scenario, intuitively, 10 matches to the bacterial sequence are less significant than 9 matches to the viral sequence.

To address these challenges, we model the read classification as a binomial experiment. We assume that the incoming DNA read is a random sequence of nucleotides, and hence the *k*-mers are random as well. We also assume that all *k*-mers from the read are independent, despite the fact that some will contain overlapping base pairs. Lastly, for both reads and references, we consider only canonical syncmers, which form a subset, *U*, of all possible 4^*k*^ *k*-mers. Given these assumptions, the probability that a random *k*-mer from the read is in the *k*-mer set *K*_*i*_ of assembly *a*_*i*_, is simply 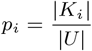. For the remainder of this section, we will refer to the canonical syncmers extracted from a read simply as *k*-mers.

When classifying a read, for each assembly *a*_*i*_ we assume a null hypothesis, *H*_*i*0_, that the matches between the read’s *k*-mers and the assembly *a*_*i*_ are not statistically significant. Then, for a given read, we enumerate all its *k*-mers, and use each enumerated *k*-mer to query our reference database. For each assembly *a*_*i*_ we maintain a counter, *x*_*i*_, that indicates the number of times a *k*-mer from the read matched to *a*_*i*_. When there are no more *k*-mers in the read, we calculate the p-value of observing *x*_*i*_ or more *k*-mer matches for each *a*_*i*_ under the assumption of a binomial experiment with statistic *P* (Bin(*n, p*_*i*_) ≥ *x*_*i*_), where *n* is the total number of *k*-mers extracted from the read being classified. Put differently, for a read with *n k*-mers, *P* (Bin(*n, p*_*i*_) ≥ *x*_*i*_) gives us the probability that we observed *x*_*i*_ or more *k*-mer matches with *a*_*i*_, given that the probability of match success for *a*_*i*_ is *p*_*i*_.

If the observed p-value for any *a*_*i*_ is lower than a predefined cutoff threshold, we reject the null hypothesis. We conclude that this is good evidence that the matching of *k*-mers between the read and the assembly *a*_*i*_ is significant, and we classify the read as coming from *a*_*i*_. If more than one null hypothesis is rejected, we choose the assembly with the lowest p-value.

We modify this procedure to account for the fact that ONT reads usually have drastically different lengths, resulting in significant variability in the number of *k*-mers, *n*, extracted from each read. Since *n* is the number of binomial trials in our tests, it can affect the comparability of the corresponding p-values. To address this, instead of considering *n* directly, we calculate our classification probability *P* based on a maximum likelihood estimate for the number of matching *k*-mers expected if exactly *n*_fixed_ *k*-mers were considered.

More precisely, we use 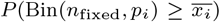 to obtain the p-value for *a*_*i*_, where 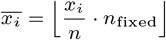 If less than *n* were queried, we do not calculate 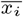 (we simply use 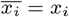), but we still use *n*_fixed_ to calculate the p-value (i.e., *P* (Bin(*n*_fixed_, *p*_*i*_) ≥ *x*_*i*_)). The reason for this is explained in Section 4. Overall, this makes a single cutoff threshold more comparable between reads of differing lengths. Moreover, because *n*_fixed_ can be relatively small, for instance we use *n*_fixed_ = 100 by default, it allows us to reduce runtime by pre-computing a p-value lookup table for all possible number of matches in the range [0, *n*_fixed_] for each assembly *a*_*i*_.

Putting this all together, the complete classification process for a single read works as follows: First, for each assembly *a*_*i*_ we pre-compute both *p*_*i*_ and the p-value lookup table. Each *p*_*i*_ is stored in the SKiM database, and the lookup table is quickly calculated at runtime using the *n*_fixed_ parameter. For each *k*-mer in the read, we use the bijection *f* to find its corresponding ARLE-compressed row in our reference database. For each *a*_*i*_ present in the row, we increment counter *x*_*i*_, and once all *n k*-mers from the read have been searched, for each *x*_*i*_ we calculate 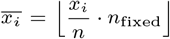 (if needed) and use 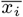 to lookup the pre-computed p-value. If the smallest p-value found is lower than the predefined cutoff threshold, we classify the read as coming from the assembly *a*_*i*_ for which we obtained that p-value. Otherwise, we report the read as unclassified. Here, the cutoff threshold is defined as 10^−*e*^, where *e* is user-defined at runtime (*e* = 12 by default).

## 3. Results

To assess the effectiveness and performance of our proposed solutions, we compared SKiM with other metagenomic classifiers under conditions that mimic adaptive sampling and real-time classification applications. In the tests, we used `Centrifuge` 1.0.4 [6], `Kraken2 2.1.3` [27], `KrakenUniq 1.0.4` [2], and CLARK 1.2.6 [18]. We also considered the more recent Taxor 0.1.3 [25], however, we found it unable to perform classification in any tests we conducted. We ran all classifiers with default parameters, except for `Centrifuge`, where we limited the number of classification results reported per read to one (using the -k 1 switch).

### 3.1 Test Data

In all tests, we ran all the tools with the same reference database. This database consisted of all archaeal, bacterial, and viral sequences from NCBI RefSeq with the annotation “Complete Genome,” downloaded together with the NCBI taxonomy on June 12, 2023. We further extended the database with 12 reference sequences for the synthetic mock microbial community from [23] that were not covered by the NCBI sequences. We refer to the resulting database as ABV.

This database represents a large, quality reference suitable for a comprehensive metagenomic classification without prior knowledge of the composition of the sequenced samples. Collectively, it spans 48,214 assemblies that take 145GB when stored in FASTA files.

As input data, we used reads from three actual ONT metagenomic sequencing experiments. The first dataset was from the GridION sequencing of a ZymoBIOMICS Microbial Community Standard with known composition, publicly available from [15] (European Read Archive accession ERR2887847). This standard contains eight species of bacteria and two species of yeast, where all bacteria have even abundance. The dataset has 3,491,390 reads and contains raw ONT signals. This enabled us to re-basecall the raw signals with `Dorado` using model dna_r9.4.1_e8_fast@v3.4 (originally, the reads were basecalled with `Guppy`, an earlier generation of ONT basecallers). We used the fast model as it is the most likely choice when performing basecalling on the computationally lightweight hardware we target. We will refer to this dataset as Even.

For the second dataset, we used MinION sequencing from [9] where the sequencing sample was a synthetic mock community of 12 bacteria with even abundance. The reads were originally basecalled with Albacore, another early generation ONT basecaller with a higher error rate than Dorado. Since the raw ONT signals were provided, we re-basecalled and demultiplexed them with `Dorado` (model dna_r9.4.1_e8_fast@v3.4) as well. The resulting dataset, which we will refer to as Bench, contains 431,287 reads.

For the third dataset, we used the MinION sequencing from [23] (NCBI Sequence Read Archive accession SRX4901586), where the sequencing sample was another synthetic mock community of 12 bacteria. More specifically, the dataset contains 10Kbp size-selected reads (i.e., only reads longer than 10Kbp), and the abundance of each bacteria is uneven, although no single species accounts for the majority of reads. The dataset has 187,507 reads basecalled with Albacore, but did not include raw ONT signals that we could re-basecall. We will refer to this dataset as BMock12-10kb.

To mimic the classification conditions of an adaptive sampling experiment, we took each of the three datasets and generated new datasets that consisted of all the same reads, but with a constrained maximum read length 𝓁. During classification, for every read longer than the maximum read length 𝓁, we considered only the first 𝓁 base pairs, and for reads shorter than 𝓁, we considered the entire read. Here, we note that the reported optimal fragment size when working with the ONT interface for adaptive sampling is around 180bp (see, e.g., [19]). Therefore, in our experiments, we consider 𝓁 of 180bp, 360bp, 720bp, and 1440bp, along with the unconstrained full read length.

### 3.2 Classification Effectiveness

To assess the effectiveness of each classifier, we used commonly accepted statistics, used by, for example, CAMI [10]. Specifically, to establish a reference point (ground truth) for the tests, we used minimap2 [10], with ONT-specific parameters, and aligned the full input reads against only those reference sequences that should be present in a sample based on its known composition. If a given read did not have a significant alignment or if there were alignments of the same quality to multiple references, we discarded it from the subsequent analysis. Otherwise, we assigned its ground truth classification to be the taxonomic identifier of the reference sequence with the most significant alignment.

We counted a classification as a true positive (TP) when its classifier-assigned identifier matched the ground truth identifier, a false positive (FP) when the classifier-assigned identifier was different from the ground truth, a false negative (FN) when the classifier did not assign any identifier, but the ground truth identifier was known, and finally, a true negative (TN) when the classifier did not assign an identifier and the ground truth identifier was not in the classifier’s reference database. We note that TNs occur only in the case of the Even dataset, since this dataset contains yeast sequences that we did not include in the ABV reference. Hence, any read classified as yeast by minimap2 would ideally remain unclassified by any classifier being tested.

To assess classification effectiveness at different taxonomic levels (i.e., species, genus), we relied on the NCBI taxonomy. This required additional care to account for sporadic situations where some species-level taxonomic identifiers have no genus-level assignment (i.e., their parent node in the taxonomic tree is labeled as “no rank”). For all such cases, we assumed that the genus-level identifier is the parent identifier of the selected species-level identifier.

We created four SKiM databases with different sub-sampling parameters to observe the impacts of these parameters on both classification and performance. When reporting results, we use `SKiM` to denote a reference database constructed with the default parameters (*k* = 15, *s* = 9 and *t* = 2), and `SKiM-k-s-t` when using other configurations. For example, SKiM-k_15_-s_13_-t_1_ refers to the configuration with *k* = 15, *s* = 13, and *t* = 1. In all SKiM classification experiments, we use the cutoff value of 10^−12^.

The results of our experiments are summarized in Table 1 and Figures 2-4. The Supplementary Information contains a complete set of raw statistics for all experiments as well.

**Table 1.** Classifier statistics for the Even dataset when 𝓁 = 360bp.

|  | Species |  |  |  | Genus |  |  |  |
| --- | --- | --- | --- | --- | --- | --- | --- | --- |
| | Recall | Precision | Accuracy | $F_1$-Score | Recall | Precision | Accuracy | $F_1$-Score |
| Centrifuge | 67.67 | 88.54 | 63.58 | 76.71 | 81.36 | 90.82 | 76.06 | 85.83 |
| Kraken2 | 63.18 | 92.49 | 61.91 | 75.08 | 86.71 | 96.63 | 84.88 | 91.40 |
| KrakenUniq | 59.22 | **93.73** | 58.95 | 72.58 | 87.41 | **97.02** | 85.81 | 91.96 |
| CLARK | 56.20 | 93.22 | 56.12 | 70.12 | 56.87 | 95.82 | 57.56 | 71.38 |
| SKiM | 80.12 | 92.66 | 76.43 | 85.93 | 80.45 | 94.65 | 77.98 | 86.98 |
| SKiM-$k_{15}$-$s_{13}$-$t_1$ | 92.22 | 92.17 | 86.12 | 92.20 | 92.37 | 94.18 | 87.90 | 93.27 |
| SKiM-$k_{15}$-$s_{14}$-$t_1$ | **94.84** | 90.09 | **86.35** | **92.40** | **94.95** | 92.13 | **88.23** | **93.52** |
| SKiM-$k_{16}$-$s_{10}$-$t_2$ | 85.59 | 93.41 | 81.57 | 89.33 | 85.82 | 95.22 | 83.06 | 90.28 |

We define Recall as $\frac{TP}{TP + FN}$ , Precision as $\frac{TP}{TP + FP}$ , and Accuracy as $\frac{TP + TN}{TP + FP + FN + TN}$ .

**Fig. 2.**
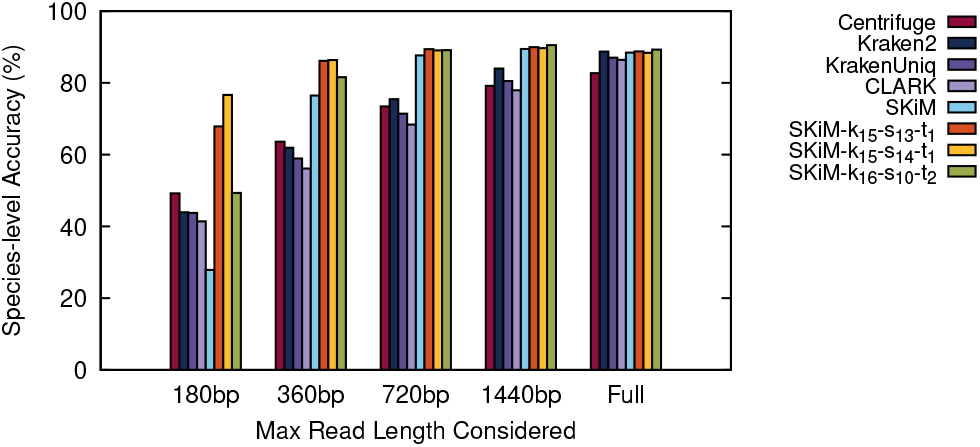
Species-level classifier accuracy on the Even reads.

**Fig. 3.**
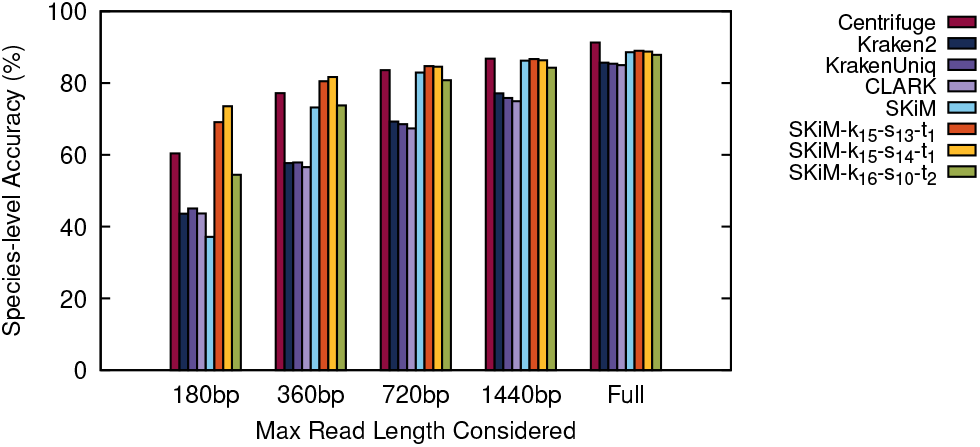
Species-level classifier accuracy on the Bench reads.

### 3.3 Performance Statistics

To assess the computational performance of the tested classifiers, we ran all tests on a server with 512GB of RAM and an AMD EPYC 7203P 8-core processor that supports 8 threads. To report the peak memory usage of each classifier when the classifier is running, we extracted the maximum resident set size (RSS) provided by the `time -v`UNIX tool. Finally, to obtain the classification throughput, we used the self-reported throughput for the classifiers that provided such information, or we calculated it based on the walltime used (excluding the time to load the reference database). These results are summarized in Table 2 and Figure 5. The throughput figures for Bench and BMock12-10kb, which are almost identical to Figure 5, are available in Supplementary Information Figures S1 and S2.

**Table 2.** Database size and memory use for each classifier on ABV.

| Classifier | Size on Disk (GB) | Peak RSS (GB) |
| --- | --- | --- |
| Centrifuge | 81.5 | 87.9 |
| Kraken2 | 62.0 | 62.9 |
| KrakenUniq | 432.7 | 430.6 |
| CLARK | 184.8 | 184.8 |
| SKiM | 12.4 | 14.6 |
| SKiM-$k_{15}$-$s_{13}$-$t_1$ | 24.1 | 28.3 |
| SKiM-$k_{15}$-$s_{14}$-$t_1$ | 35.2 | 40.2 |
| SKiM-$k_{16}$-$s_{10}$-$t_2$ | 17.9 | 26.5 |

**Fig. 4.**
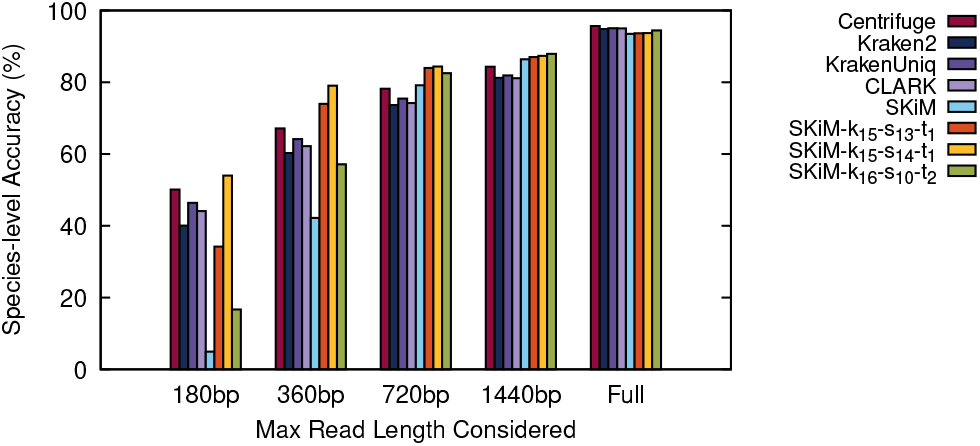
Species-level classifier accuracy on the BMock12-10kb reads.

**Fig. 5.**
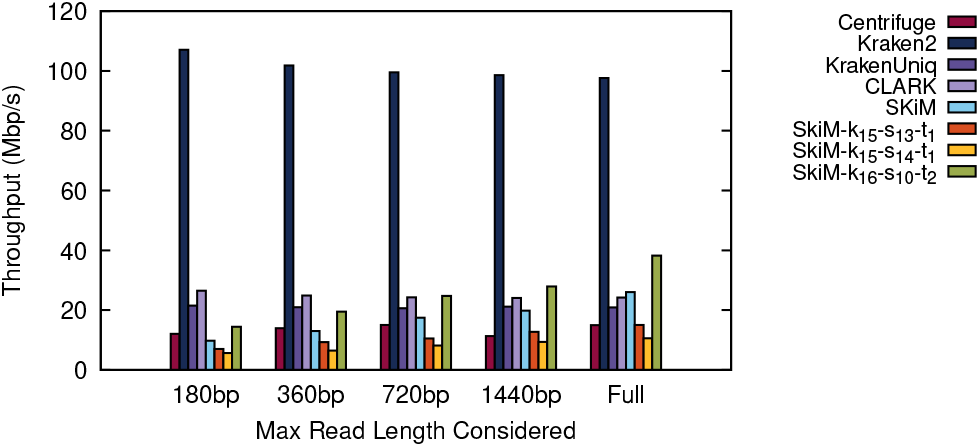
Classifier throughput on the Even reads.

To understand the performance of our proposed compression method, we created variants of the default SKiM database using: a random column ordering of input assemblies and the NRLE compression scheme (denoted as `Random NRLE`), a random column ordering and our ARLE compression scheme (denoted as `Random ARLE`), the TSP-heuristic induced column ordering and NRLE scheme (denoted as `SKiM NRLE`), and the TSP-heuristic ordering with ARLE scheme (which is simply referred to as `SKiM`). We additionally include SKiM databases constructed with different *k*-mer and sub-sampling parameters. As previously described, these variants are labeled as SKiM-k-s-t. The properties of each resulting database are summarized in Table 3. For the Random NRLE and Random ARLE tests, we report the average over five different random orderings.

**Table 3.**
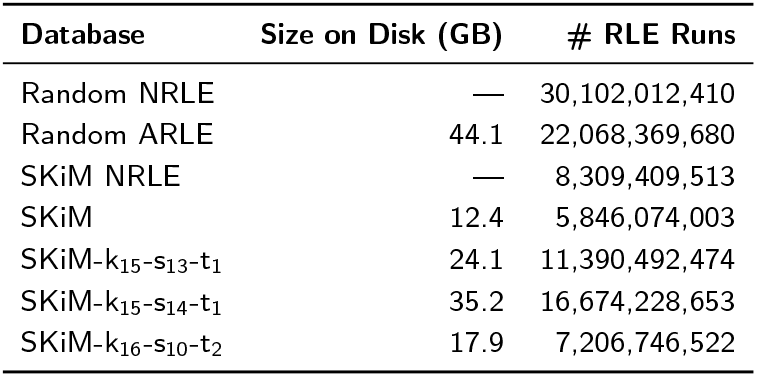
Effects of compression on the size of the SKiM ABV database.

## 4. Discussion

SKiM outperforms the tested classifiers in terms of recall and accuracy when limited to the first 720bp of a read while maintaining comparable precision. Moreover, its classification performance when considering at least 720bp is comparable to that of the entire read, which is not the case for other tools. Furthermore, for Even and Bench datasets, which were basecalled using Dorado, SKiM achieves similar performance when considering only 360bp. Table 1 shows that when limited to 360bp on the Even dataset, SKiM has the highest recall, accuracy, and *F*_1_-Score of all classifiers, while the precision is comparable to that of other tools. In the context of adaptive sampling, this allows SKiM to accurately classify reads earlier than other tools. This property is critical as it means that SKiM can correctly detect reads that are not of interest earlier and can be ejected by a sequencer, thus freeing resources.

However, SKiM is not without limitations. From Figures 2-4 we can see that SKiM with default parameters has significantly lower accuracy when reads are restricted to the first 180bp. This is caused by a larger number of false negative assignments (see raw data in Supplementary Information), suggesting that when the read length is too short we have insufficient information to accurately estimate 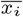. More specifically, with the default sub-sampling parameters, only 12% of all possible *k*-mers are canonical syncmers. That means that a 180bp read would only provide approximately 21 *k*-mer queries to calculate 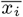. Because this is below the default *n*_fixed_ = 100, we do not perform any scaling on the number of matches *x*_*i*_, but we still calculate p-values based on *n*_fixed_ (recall Section 2.2). We could instead calculate the p-value based on the true number of queries *n*, use a larger cutoff value, or always calculate 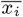. Indeed, we observed that each possible solution decreased the number of false negatives, increased the number of true positives, and increased the overall accuracy to outperform all other classifiers. But, it also introduced a significant number of false positives. Within the context of adaptive sampling, false positive decisions are more problematic than false negatives. If a read is falsely classified, it may be incorrectly ejected before more base pairs are sequenced, allowing a better classification to be made.

SKiM performance on the Bmock12-10kb dataset (Figure 4) is slightly worse compared to the other two test datasets. This dataset is the only dataset basecalled with Albacore, a very early ONT basecaller that has a high error rate compared to the newest basecaller Dorado. Consequently, we attribute SKiM’s worse performance on this dataset to the lower quality of the reads.

Table 2 shows that SKiM runs with a fraction of the memory, as low as 14.6GB for all of ABV, while the next best classifier, `Kraken2`, requires 62.9GB. As Table 3 shows, the proposed lossy compression methods play a crucial role in SKiM’s database sizes. More specifically, it shows that a random column ordering of ABV requires around four times as many runs as the nearest neighbor heuristic ordering (for both NRLE and ARLE). It also shows that the number of runs required by the ARLE representations is around 70% of the corresponding NRLE representation (for both the random and TSP-heuristic ordering). Without these optimizations, SKiM’s default parameter database would be comparable in size to `Kraken2`.

While the TSP-heuristic ordering provides significant compression, it implicitly requires that we compute the distance between each pair of assembly bitmaps (*K*_*i*_, *K*_*j*_). For large assembly sets, this computation may become a bottleneck in database construction. However, this can be addressed by augmenting the TSP-heuristic with taxonomic information. Specifically, we can partition the input assemblies by the species, genus, or family to which they belong, and find the TSP-heuristic ordering on each partition separately. Then, the final ordering can be obtained by linking partitions together, using the smallest distances between partitions. This approach avoids the quadratic runtime of computing pairwise distances between all pairs of assemblies.

Lastly, Figure 5 shows that SKiM has a classification throughput of 10-25Mbp/s, which is comparable to other tools, and is outperformed only by `Kraken2.` For comparison, at maximum theoretical throughput, a MinION flow cell can concurrently sequence 512 molecules at 450bp/s each [26], for a total of 230Kbp/s. While hardware at the point of sequencing may not be as fast or have as many available cores as the processor used in our experiments, SKiM’s throughput is still comfortably faster than the maximum theoretical sequencing throughput. This is true even for GridION devices, capable of sequencing on five MinION flow cells concurrently. Consequently, we would not expect SKiM to cause a bottleneck during adaptive sampling or real-time classification.

### 4.1 Implications for Pangenomes and Strain-level Classification

Reference databases used in metagenomics frequently involve highly similar sequences, for example, coming from multiple strains of the same bacterial species. When these sequences are treated as the same classification target, as in the case for tools like `CLARK, Kraken2, Centrifuge`, etc., significant index reduction can be achieved. However, this design does not allow for classification at the level of reference assembly or sequence. Because SKiM indexes each reference assembly individually, it may be better equipped for pangenome and strain-level classification experiments. Furthermore, highly similar assemblies are likely to be significantly compressed in a SKiM database but without information loss.

As a proof of concept, we used SKiM with default parameters to index all DNA sequences for SARS-CoV-2 deposited in the NCBI SARS-CoV-2 Data Hub [30] throughout 2020. With a total of 47,160 sequences (about 1.4GB when stored as FASTA files), SKiM was able to create an index of just 5MB. This is a promising result. For example, such a 5MB index could simply be added to the existing ABV database with minimal additional overhead, allowing for more robust classification of many SARS-CoV-2 variants.

## 5. Competing interests

No competing interest is declared.

## 6. Author contributions statement

T.S. and J.Z. developed the methods and designed experiments, T.S. implemented the methods and conducted the experiments, T.S. and J.Z. analyzed the results. T.S. and J.Z. wrote and reviewed the manuscript.

## 7. Acknowledgments

This work is part of the SMARTEn Project [29] and is supported in part by funds from the National Science Foundation (CNS-1910193).

## 8. Data Availability

SKiM is available at https://gitlab.com/SCoRe-Group/SKiM. All datasets were derived from sources in the public domain: Even [15] (https://www.ebi.ac.uk/ena/browser/view/ERR2887847), Bench [9] (https://zenodo.org/records/3600229), and BMock12-10kb [23] (https://www.ncbi.nlm.nih.gov/sra/SRX4901586).

## Notes

### Competing Interest Statement

The authors have declared no competing interest.

## References

1. K.S. Booth and G.S. Lueker. Testing for the consecutive ones property, interval graphs, and graph planarity using PQ-tree algorithms. Journal of Computer and System Sciences, 13(3):335–379, 1976.

2. F.P. Breitwieser, D.N. Baker, and S.L. Salzberg. KrakenUniq: confident and fast metagenomics classification using unique k-mer counts. Genome Biology, 19(1):198, 2018.

3. S. Chambi, D. Lemire, O. Kaser, and R. Godin. Better bitmap performance with Roaring bitmaps. Software: Practice and Experience, 46(5):709–719, 2016.

4. R. Edgar. Syncmers are more sensitive than minimizers for selecting conserved k-mers in biological sequences. PeerJ, 9:e10805, 2021.

5. D. Johnson, S. Krishnan, J. Chhugani, S. Kumar, and S. Venkatasubramanian. Compressing large boolean matrices using reordering techniques. In International Conference on Very Large Data Bases, VLDB ‘04, pages 13–23, Toronto, Canada, 2004. VLDB Endowment.

6. D. Kim, L. Song, F.P. Breitwieser, and S.L. Salzberg. Centrifuge: rapid and sensitive classification of metagenomic sequences. Genome Research, 26(12):1721–1729, 2016.

7. E.J. Kipp, L.L. Lindsey, B. Khoo, et al. Metagenomic surveillance for bacterial tick-borne pathogens using nanopore adaptive sampling. Scientific Reports, 13(1):10991, 2023.

8. S.Y. Ko, L. Sassoubre, and J. Zola. Applications and Challenges of Real-time Mobile DNA Analysis. In International Workshop on Mobile Computing Systems & Applications, pages 1–6, Tempe Arizona USA, 2018. ACM.

9. R.M. Leidenfrost, D. Pöther, U. Jäckel, and R. Wünschiers. Benchmarking the MinION: Evaluating long reads for microbial profiling. Scientific Reports, 10(1):5125, 2020. Publisher: Nature Publishing Group.

10. H. Li. Minimap2: pairwise alignment for nucleotide sequences. Bioinformatics, 34(18):3094–3100, 2018.

11. M. Loose, S. Malla, and M. Stout. Real-time selective sequencing using nanopore technology. Nature Methods, 13(9):751–754, 2016.

12. S. Martin, D. Heavens, Y. Lan, et al. Nanopore adaptive sampling: a tool for enrichment of low abundance species in metagenomic samples. Genome Biology, 23(1):11, 2022.

13. A.J. Mikalsen and J. Zola. Coriolis: enabling metagenomic classification on lightweight mobile devices. Bioinformatics, 39(Supplement 1):i66–i75, 2023.

14. G. Navarro. Compact Data Structures: A Practical Approach. Cambridge University Press, 2016.

15. S.M. Nicholls, J.C Quick, S. Tang, and N.J. Loman. Ultra-deep, long-read nanopore sequencing of mock microbial community standards. GigaScience, 8(5):giz043, 2019.

16. N.A. O’Leary, M.W. Wright, J.R. Brister, et al. Reference sequence (RefSeq) database at NCBI: current status, taxonomic expansion, and functional annotation. Nucleic Acids Research, 44(D1):D733–D745, January 2016.

17. F. Olken and D. Rotem. Rearranging Data to Maximize the Efficiency of Compression. In ACM SIGACT-SIGMOD Symposium on Principles of Database Systems, 1985.

18. R. Ounit, S. Wanamaker, T.J. Close, and S. Lonardi. CLARK: fast and accurate classification of metagenomic and genomic sequences using discriminative k-mers. BMC Genomics, 16(1):236, 2015.

19. A. Payne, N. Holmes, T. Clarke, et al. Readfish enables targeted nanopore sequencing of gigabase-sized genomes. Nature Biotechnology, 39(4):442–450, 2021.

20. J. Quick, N.J. Loman, S. Duraffour, et al. Real-time, portable genome sequencing for Ebola surveillance. Nature, 530(7589):228–232, 2016. Publisher: Nature Publishing Group.

21. M. Roberts, W. Hayes, B.R. Hunt, S.M. Mount, and J.A. Yorke. Reducing storage requirements for biological sequence comparison. Bioinformatics, 20(18):3363–3369, 2004.

22. A. Schleimer, D.S. Wilkerson, and A. Aiken. Winnowing: local algorithms for document fingerprinting. In ACM SIGMOD International Conference on Management of Data, SIGMOD ‘03, pages 76–85, New York, NY, USA, 2003. Association for Computing Machinery.

23. V. Sevim, J. Lee, R. Egan, et al. Shotgun metagenome data of a defined mock community using Oxford Nanopore, PacBio and Illumina technologies. Scientific Data, 6(1):285, 2019.

24. J. Ulrich, L. Epping, T. Pilz, et al. Nanopore adaptive sampling effectively enriches bacterial plasmids. mSystems, 9(3):e00945– 23, 2024.

25. J. Ulrich and B.Y. Renard. Fast and space-efficient taxonomic classification of long reads with hierarchical interleaved XOR filters. Genome Research, 34(6):914–924, 2024.

26. Y. Wang, Y. Zhao, A. Bollas, Y. Wang, and K.F. Au. Nanopore sequencing technology, bioinformatics and applications. Nature Biotechnology, 39(11):1348–1365, 2021.

27. D.E. Wood, J. Lu, and B. Langmead. Improved metagenomic analysis with Kraken 2. Genome Biology, 20(1):257, 2019.

28. D.C. Wrenn and D.M. Drown. Nanopore adaptive sampling enriches for antimicrobial resistance genes in microbial communities. GigaByte, 2023:gigabyte103, 2023.

29. SMARTEn, 2025. http://score-group.org/?id=smarten.

30. NCBI SARS-CoV-2 Data Hub, 2025. https://www.ncbi.nlm.nih.gov/labs/virus/vssi/.

