## Supplementary Text for "SKiM: Accurately Classifying Metagenomic ONT Reads in Limited Memory"

### Supplementary Information: SKiM

#### 1 Theorem Proof

**Lemma 1.** *Given an arbitrary bit string $B$, Algorithm 1 encodes $B$ in the fewest possible number of runs (run of zeros, run of ones, or uncompressed).*

*Proof.* By "greedy stays ahead."

We start by assuming that the maximum length of a run block is $2^{w-2} - 1$ and an uncompressed block holds $w - 1$ bits of the original string.

Let $G$ be the list of runs (zeros, ones, or uncompressed) produced by running Algorithm 1 (the greedy algorithm) on $B$ and let $\mathcal{O}$ be the runs of an optimal solution, that, by definition, encodes $B$ in the fewest possible runs.

Considering the first $x$ runs of a solution, our progress measure is how many bits of the original string $B$ are encoded by these $x$ runs. Let $g_i$ be the $i^{th}$ progress measure from $G$, e.g., $g_3$ is how many bits of $B$ are encoded by the first three runs. Likewise, let $o_j$ is the $j^{th}$ progress measure of $\mathcal{O}$.

We want to prove that, for all $k$, $g_k \geq o_k$. In other words, that the first $k$ runs of $G$ encode at least as many bits as the first $k$ runs of $\mathcal{O}$ for all values of $k$.

We assume that, w.l.o.g., runs of zeros or ones can be considered the same case, which we refer to as compressed runs. Also, for the simplicity of the proof, we assume that compressed runs will not overflow, although we could extend the proof to not have this assumption. Finally, we point out two key invariants of the greedy algorithm: First, assuming the first $x$ runs encode the first $\ell$ bits of $B$, then adding the next run $(x + 1)$ will always encode at least the first $\ell + w - 1$ bits of $B$ (adding an uncompressed run), where $w$ is the word size. In other words, the greedy algorithm will never choose to encode less than $w - 1$ bits in the next run. Second, if there is a sequence of the same bit, starting at position $\ell + 1$, that extends beyond position $\ell + w - 1$, then the greedy algorithm will choose to encode the next run as a compressed run that is as long as possible. More precisely, if the first bit that is opposite of $B[\ell + 1]$, starting from position $\ell + 1$, occurs at position $n$, and $n - 1 > \ell + w - 1$, the greedy algorithm will encode out to position $n - 1$ using a compressed run next.

We will now prove the original statement, by induction:

*Base case.* There are two options for the optimal solution on the first run. First, the optimal solution could use an uncompressed run and encode the first $w - 1$ bits of $B$. As stated by the first invariant, the greedy algorithm will always encode at least this many bits from the starting position. Otherwise, the optimal solution uses a compressed run out to some position $m$. If $m \leq w - 1$, then the greedy algorithm will still encode at least as many bits by the first invariant. If $m > w - 1$, then there must be some position, $n$, at which the first bit that is opposite of $B[0]$ occurs, noting that $m \leq n - 1$. Because $m > w - 1$, then $n - 1 > w - 1$ must be true, and the greedy algorithm would choose to encode out to position $n - 1$ using a compressed run according to the second invariant. In all cases, $g_1 \geq o_1$.

*Inductive Step.* Assume that $g_k \geq o_k$. We want to prove that $g_{k+1} \geq o_{k+1}$. This case is very similar to the base case, with the difference being that the greedy algorithm *could* start with more of $B$ encoded. If the optimal solution uses an uncompressed run, $o_{k+1} = o_k + w - 1$. From our first invariant, we know that $g_{k+1} \geq g_k + w - 1$, so $g_{k+1} \geq o_{k+1}$ is true. If the optimal solution uses a compressed run, its total encoding will be out to some position $o_{k+1} = m$. If $m \leq g_k + w - 1$, then $g_{k+1} \geq o_{k+1}$ is still true by the first invariant. If $m > g_k + w - 1$, then there must be some position, $n$, where, starting at position $o_k + 1$, the first bit opposite of $B[o_k + 1]$ occurs, again noting that $m \leq n - 1$. Because $m > g_k + w - 1$, then $n - 1 > g_k + w - 1$ must be true, and the greedy algorithm would choose to encode out to position $n - 1$, starting from position $g_k$, using a compressed run according to the second invariant. In all cases, $g_{k+1} \geq o_{k+1}$.

Based on this proof by induction, we assert that for every run in the optimal solution, the greedy solution encodes at least as many bits from the original bit string $B$, maybe more, in the same number of runs. Therefore, by the end of the greedy encoding, the total number of runs required by the greedy algorithm must be less than or equal to the number of runs required by an optimal solution (or simply equal to, because $\mathcal{O}$ was assumed to be optimal). □

**Theorem 1.** *Let* ADAPTIVE-BLOCKS$(M)$ *be the total number of runs required to represent any bit matrix $M$, obtained by running Algorithm 1 on each of the rows of $M$. Then* ADAPTIVE-BLOCKS$(M) \leq$ NAIVE-RUNS$(M)$ *for any bit matrix $M$.*

*Proof.* First, we assume that the maximum length of a run block is $2^{w-2} - 1$ for both ADAPTIVE-BLOCKS$(M)$ and NAIVE-RUNS$(M)$ and an uncompressed block holds $w-1$ bits of the original string for ADAPTIVE-BLOCKS$(M)$. By Lemma 1, Algorithm 1 encodes an arbitrary bit string $B$ in the fewest possible blocks. In other words, running Algorithm 1 on each row in $M$ results in each row requiring the fewest number of possible blocks given the constraints. Therefore, by definition, each row $j$'s ARLE representation requires a number of blocks less than or equal to $j$'s NRLE representation. Since this can be said for every row in the matrix, we conclude that ADAPTIVE-BLOCKS$(M) \leq$ NAIVE-RUNS$(M)$ for any bit matrix $M$. □

### 2 Additional Tables and Figures

Throughput graphs for the Bench and Bmock12-10kb reads are shown in Figure S1 and Figure S2 respectively. Tables containing raw numbers to calculate recall, precision, accuracy, etc. for all experiments are available in the supplementary Excel files.

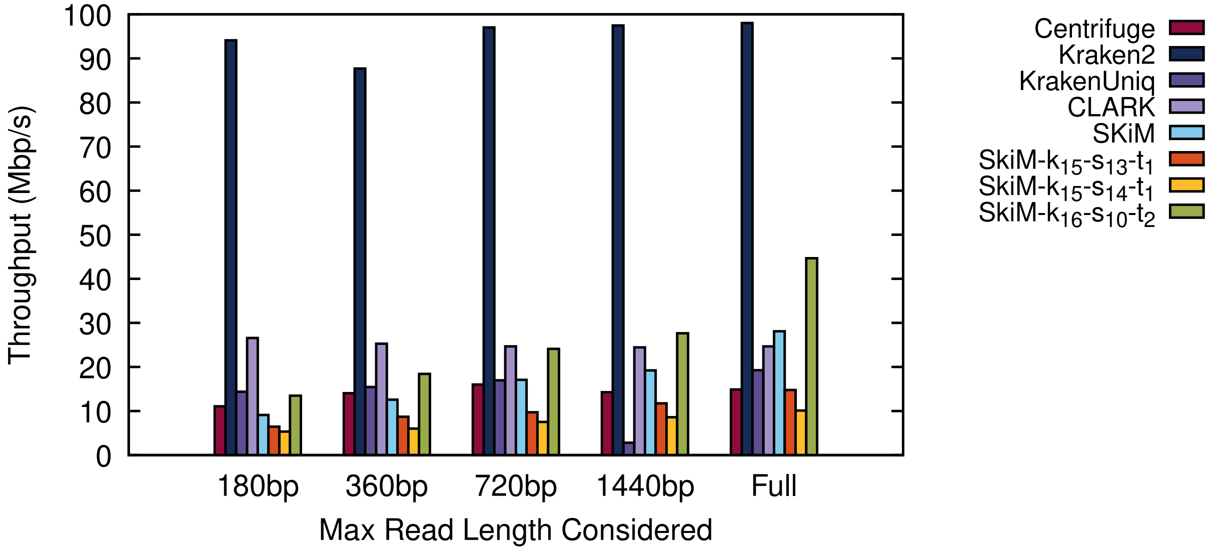


Figure S1: Classifier throughput on the Bench reads.

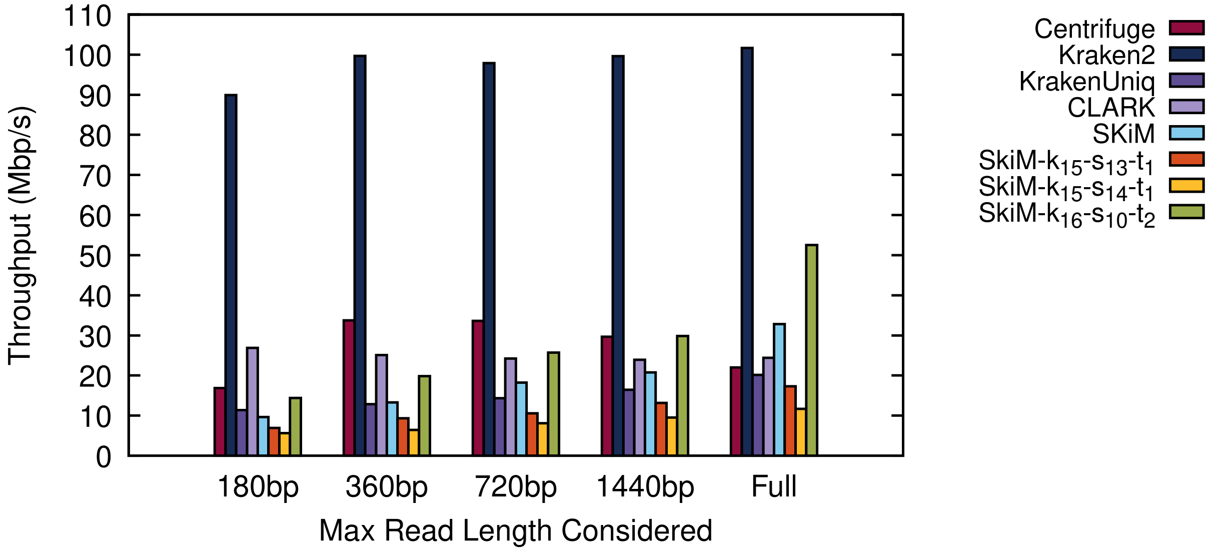


Figure S2: Classifier throughput on the BMock12-10kb reads.
